# The enigmatic role of fungal annexins: the case of *Cryptococcus neoformans*

**DOI:** 10.1101/536193

**Authors:** Maria Maryam, Man Shun Fu, Alexandre Alanio, Emma Camacho, Diego S. Goncalves, Eden E. Faneuff, Nina T. Grossman, Arturo Casadevall, Carolina Coelho

**Author notes:** Contributed equally for this work (Man Shun Fu, Alexandre Alanio). Contributed equally for this work (Arturo Casadevall, Carolina Coelho). To whom correspondence should be addressed: Carolina Coelho, PhD. Current Affiliation: Medical Research Council Centre for Medical Mycology, Institute of Medical Sciences, University of Aberdeen, Ashgrove Road West, Aberdeen, UK AB25 2ZD and College of Life and Environmental Sciences, University of Exeter, Stocker Road, Exeter, UK EX4 4QD.

## Abstract

Annexins are multifunctional proteins that bind to phospholipid membranes in a calcium-dependent manner. Annexins play a myriad of critical and well-characterized roles in mammals, ranging from membrane repair to vesicular secretion. The role of annexins in the kingdoms of bacteria, protozoa and fungi have been largely overlooked. The fact that there is no known homologue of annexins in the model organism may contribute to this gap in knowledge. However, annexins are found in most medically important fungal pathogens, with the notable exception of *Candida albicans.* In this study we evaluated the function of the one annexin gene in *Cryptococcus neoformans*, a causative agent of cryptococcosis. This gene CNAG_02415, is annotated in the *C. neoformans* genome as a target of calcineurin through its transcription factor Crz1, and we propose to update its name to cryptococcal annexin, AnnexinC1. *C. neoformans* strains deleted for AnnexinC1 revealed no difference in survival after exposure to various chemical stressor relative the wild type, as well as no major alteration in virulence or mating. The only alteration observed in strains deleted for AnnexinC1 was a small increase in the titan cells formation *in vitro*. The preservation of annexins in many different fungal species suggests an important function, and therefore the lack of a strong phenotype for annexin-deficient *C. neoformans* is suggestive of either redundant genes that can compensate for the absence of AnnexinC1 function or novel functions not revealed by standard assays of cell function and pathogenicity.

**Importance:** Cryptococcus neoformans is the deadliest human fungal pathogen, causing almost 200,000 deaths each year. Treatment of this lethal infection is lengthy, and in some patients therapy is not curative and patients require lifelong therapy. Fundamental research in this yeast is needed so that we can understand mechanisms of infection and disease and ultimately devise better therapies. In this work we investigated a fungal representative of the annexin family of proteins, specifically in the context of virulence and mating. We find that the cryptococcal annexin does not seem to be involved in virulence or mating but affects generation of titan cells, enlarged yeast cells that are detected in the lungs of mammalian hosts. Our data provides new knowledge in an unexplored area of fungal biology.

## Introduction

Annexins are a multifunctional family of proteins that bind to phospholipid membranes in a calcium-dependent manner. The bridging of membranes by annexins is calcium dependent and is mediated through a structural domain consisting of about 70 amino acids in consecutive alpha helical conformations known as the ‘annexin fold’ (1).

Annexins have been reported in all domains of life, except Archaebacteria (2, 3). In plants, annexins are important for immunity and resistance to nutrient stress (4). In mammals, annexins play a critical role in membrane signaling, encompassing plasma membrane repair, gene regulation, organelle trafficking, endosomal fusion, endocytosis and exocytosis. Therefore, annexins are indispensable for tissue functions such as clotting, immunity, among others (1, 5-7). In mammals, annexins are found in the nucleus, the cytosol, and cell surface, and are also secreted into the extracellular milieu.

A recent review revealed a paucity of studies about microbial annexins, but the available information suggests that these proteins are critical for the virulence of certain pathogens (8, 9). Giardin from *Giardia* (10) is an immunodominant annexin protein (11). Administration of giardin to mice induced protective immunity against *Giardia* and is under investigation for its vaccine potential (8). The *Burkholderia* JOSHI_001 protein possesses both a colicinD toxin domain and an annexin domain (9), which reinforces the notion that microbial annexins contribute to microbial virulence.

Very little information is available on fungal annexins. In *Neurospora crassa*, no phenotype was found during phenotype mutation efforts (12). The absence of an annexin gene in *Saccharomyces cerevisiae* and *Candida* spp. may have contributed to the lack of information on fungal annexins, given that these organisms are the most extensively studied fungi at the cellular level. However, annexin genes can be found in all basidiomycota and the majority of medically important ascomycota. Thus far, annexin deletion mutants were characterized in *Aspergillus* (13-16), a genus with 3 recognizable annexins, and *Thermomyces lanuginosus* (17). Deletion of *anxc1* in *A. niger* or *anxc3* in *A. fumigatus* did not result in growth or protein secretion defects (13). Deletion of *anxc3* (previously *anxc4*) of *A. fumigatus* was associated with very subtle changes in the protein secretion profile (13). In *T. lanuginosus*, annexinC7 is involved in conidia formation and resistance to oxidative stress (17). In amoeba *Diciostelyium discoideum*, one annexin gene is found and its deletion led to a delay in growth in low Ca^2+^ conditions (18).

*Cryptococcus neoformans* is the causative agent of cryptococcosis, a life-threatening infection that often results in high mortality and morbidity (19). The cryptococcal genome contains one predicted annexin gene, CNAG_02415, annotated as AnnexinXIV. A literature search for references to AnnexinXIV, fungal annexin or CNAG_02415 found no reports on the function of this gene. A genetic screen showed that CNAG_02415 deletion led to altered susceptibility to a range of small molecule compounds (20). Further, CNAG_02415 is a target of Crz1, an effector of the calcineurin pathway which is essential for virulence in *C. neoformans* (20-22). Chip-Sequencing data identified AnnexinC1 as a target of Crz1 (23), a major calcineurin-dependent transcription factor (22). This finding posits a role for AnnexinC1 as a component of the Crz1-calcineurin response, involved in high-temperature growth, cell wall stability and heavy metal susceptibility (22, 24). Available gene and protein expression datasets in *C. neoformans* detected no alterations in expression of AnxC1 expression upon infection (25), nor was it was detected as secreted into the extracellular milieu (26, 27). In a transcriptomic dataset, absence of Rim101 led to an increase in expression of *ANXC1* when placed for 3h in Dulbecco’s Modified Eagle Medium (28, 29), which suggest *ANXC1* is additionally regulated by Rim101 (28), an important cell morphology regulator. Based on its regulation by two important fungal signaling pathways, we decided to characterize the function of this gene. We posited that annexin in *C. neoformans* could play a role in fungal pathogenesis and its life cycle. We found that annexin deletion affected titan cell production but had no contribution to virulence in a mouse model. As with in the other fungi cited above, the molecular function of the cryptococcal annexin remains cryptic, but we provide a first step in characterizing its functions in the fungal world.

## Results

### Genetic information and previous data

Annexins share a common structural feature known as the annexin domain, a fourfold repeat of alpha helixes made up of about 70 amino acids. This domain binds Ca^2+^, which in turn mediates phospholipid binding. Despite this common structural feature of the annexin fold, annexins show significant diversity between kingdoms (9,29). Phylogenetic analysis shows that Annexins can be grouped in 5 clades (A through E). Animal annexins manifest highly conserved motifs whereas annexins in plants show motif diversity (3). In the kingdom Protista there is even greater diversity among annexins and these organisms generally express more annexins genes than in animal genomes (10). Bacterial annexin sequences are so diverse that the existence of non-classical annexins was postulated (9). Mammals and Protista have multiple different annexins in their genomes, but only one annexin has been identified in the cryptococcal genome, the gene product of CNAG_02415 (13, 16). We performed homology gene searches using DNA and protein sequences of mammalian (*Mus musculus*), plant (*Arabidopsis thaliana)*, fungal (*Aspergillus* spp.) and bacterial ‘non-classical’ annexins (9) against the cryptococcal genome to identify other annexin-like genes, but found no other homologue. Thus, we concluded that CNAG_02415 is the only classical annexin in *C. neoformans*. We extracted the predicted protein sequences of annexins from the fungal kingdom and built a phylogenetic tree (Figure 1). Fungal annexins are phylogenetically closer to *Dictyostelium discoideum* amoeba model organism than to human or plant annexins, and consequently, fungal and amoeba annexins are grouped in clade C. The sequence of fungal annexins differs significantly from that of their mammalian and plant counterparts (16), with the 3^rd^ and 4^th^ folds of fungal annexins harboring unusual sequences (30) while still retaining clear and recognizable annexin folds. CNAG_02415 encodes a protein with four of the characteristic annexin domains, one predicted calcium binding site (Figure 1) on the C-terminus, and a NH_2_ terminal head (1). The annotation of the tau domain of DNA polymerase III at the N-terminus is cryptic but may be related to nucleotide-binding properties, which have been reported for annexins (31). This cryptococcal gene is annotated as AnnexinXIV, likely due to homology with the first described fungal annexin in *Neurospora crassa* (18). Based on the nomenclature conventions of annexins (32), we name the gene product as annexinC1 (AnxC1) and the gene *ANXC1*.

**Figure 1.**
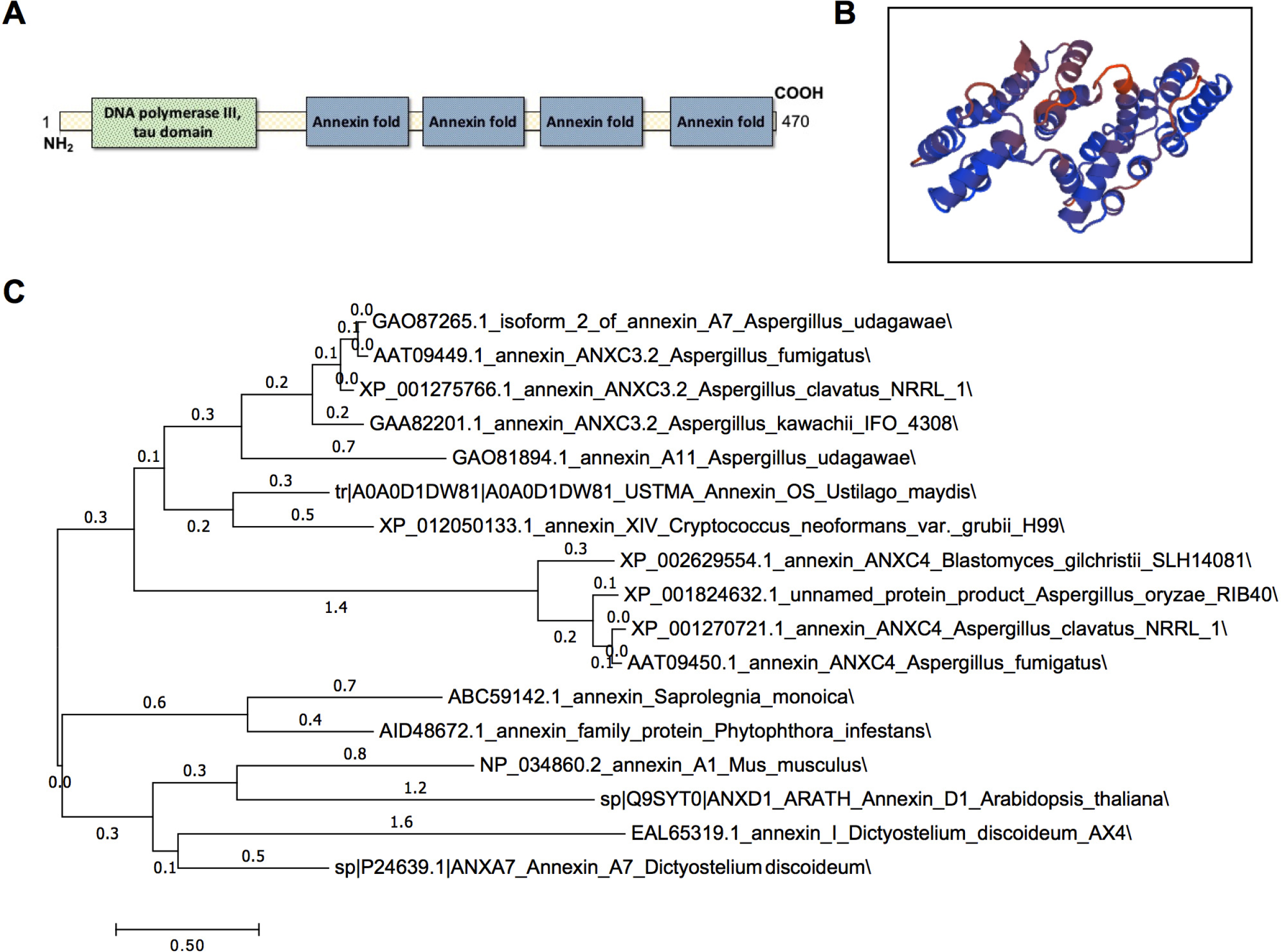
Predicted structure and phylogeny of cryptococcal annexin. A) Diagram of location of conserved domains (as predicted by NCBI Conserved Domains); Prediction of annexin C1 structure (J9VS56) as described in Materials and Methods. C) Molecular Phylogenetic analysis by Maximum Likelihood method. The evolutionary history was inferred by using the Maximum Likelihood method based on the Le_Gascuel_2008 model [43]. The tree with the highest log likelihood (−7069.32) is shown. Initial tree(s) for the heuristic search were obtained automatically by applying Neighbor-Join and BioNJ algorithms to a matrix of pairwise distances estimated using a JTT model, and then selecting the topology with superior log likelihood value. A discrete Gamma distribution was used to model evolutionary rate differences among sites (5 categories (+G, parameter = 2.7919)). The tree is unrooted with numbers indicating number of substitutions per site. The analysis involved 17 amino acid sequences. All positions with less than 95% site coverage were eliminated. That is, fewer than 5% alignment gaps, missing data, and ambiguous bases were allowed at any position. There were a total of 250 positions in the final dataset. Evolutionary analyses were conducted in MEGA7 [44].

### Generation of a new deletion mutant

We constructed AnxC1-deficient strains in two independent rounds of biolistic transformation using a H99 parental strain (lineage E, as indicated in Materials and Methods. After individual colony screening we generated 10 isolates indistinguishable by PCR and Southern blot (Supplementary Figure 1). We randomly selected 4 of these clones *AnxC1*-deleted (*anxC1*Δ) and their corresponding parental strain (wild-type H99) for subsequent studies.

### Functions of cryptococcal AnnexinC1

AnnexinC1 is reported to be a target of Crz1, a major component of the calcineurin pathway (22). To ascertain possible defects in responses dependent on the calcineurin pathway, we performed several phenotypic tests in functions strongly associated with the calcineurin pathway (Figure 2). We found no alteration in growth when *anxC1*Δ strains were exposed to the calcineurin inhibitor FK506, high temperature, or low calcium conditions due to addition of calcium chelator (1,2-bis(o-aminophenoxy)ethane-N,N,N′,N′-tetraacetic acid (BAPTA). The calcineurin pathway is also involved in defense against cell wall stress, caused by hyperosmotic stress, calcofluor white (CFW), congo red or caffeine. We found no evidence for a defect in cell wall stress for *anxC1*Δ compared to wild-type when exposed to either of these compounds. Other chemical and physical stresses, such as pH, heavy metals, UV damage were also tested and no alteration in susceptibility was detected (22, 33). In previous work, *anxC1*Δ deletion strains in an H99 background showed a small increase in susceptibility to cell wall stress and other chemical stressors (20, 22). We had access to some of these strains as they originate from an available deletion library in *C. neoformans*, in an H99 background (34). We had also access to a bigger deletion library in a KN99 background (from Hiten Madhani laboratory). To confirm the lack of cell wall stress alterations in the mutants we generated and ascertained the effect of AnxC1 in a different strain of *C. neoformans* we tested the deletion mutants from the available libraries for FK506 susceptibility, high temperature and the cell-wall stressor CFW. As previously reported we found that the *anxC1*Δ in the H99 background had a small increase in susceptibility to FK506 and high temperature (39°C) (22). These defects were not observed in the *anxC1*Δ from the KN99 background (Supplemental Figure 2). To test the contribution of *anxC1*Δ to a wider range of calcineurin dependent conditions we measured sexual mating and filamentation efficiency. We detected similar pattern and timing in mating when we crossed our *anxC1*Δ strains (α mating type) with an isolate from **a** mating type. Finally, we tested susceptibility to fluconazole and amphotericin B and determined an MIC was 2 µg/ml and 0.5-1 µg/ml for all three experiments, respectively, and no difference in MIC between wild-type and deletion strains (Figure 3).

**Figure 2.**
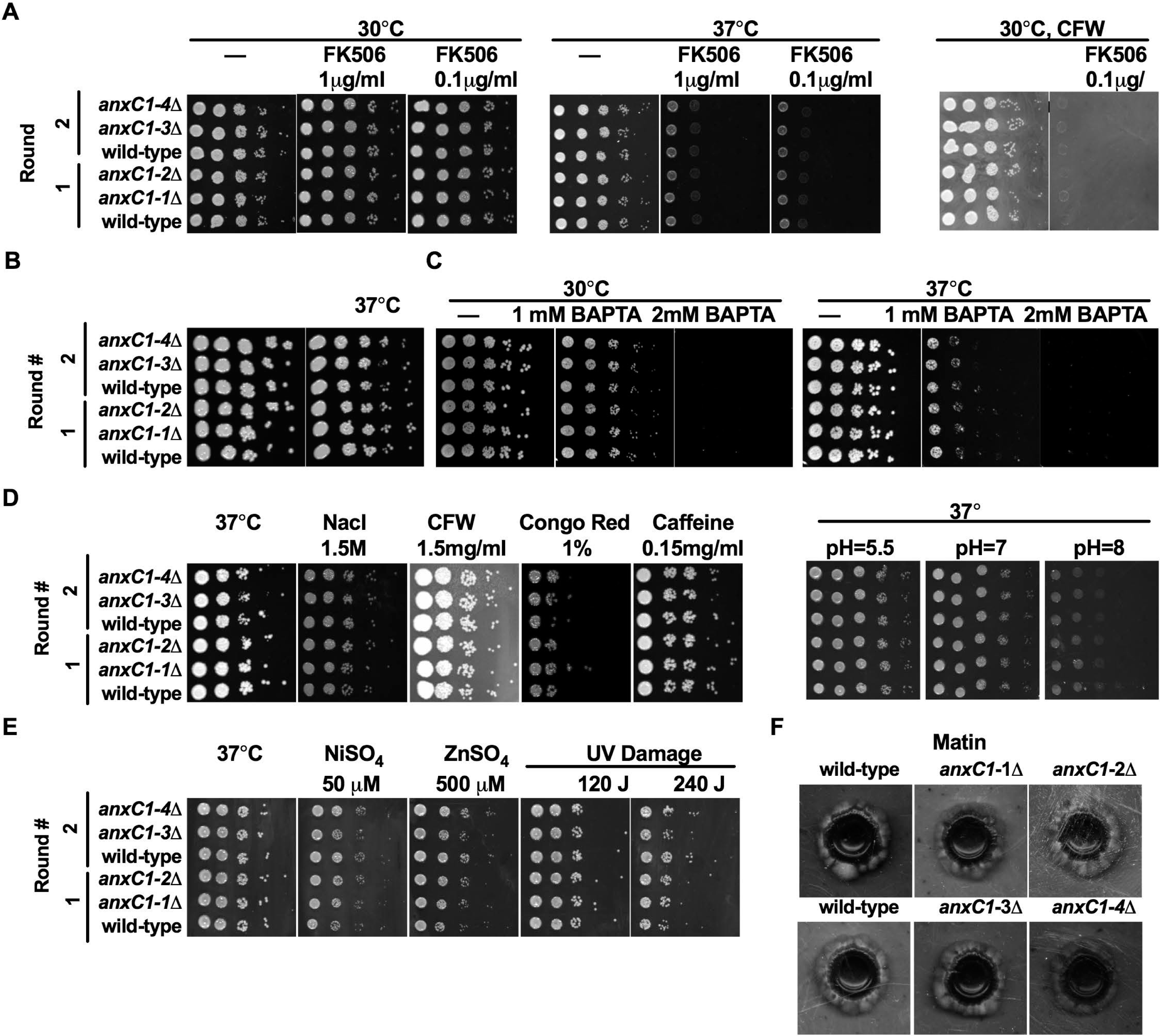
Deletion of AnnexinC1 does not affect phenotypes associated with calcineurin. AnnexinC1 deletion mutants are not susceptible to A) calcineurin inhibitors; B) high temperature; C) low calcium concentrations due to BAPTA-chelation of calcium; D) cell wall stress or alkaline pH stress; E) nickel stress or UV damage; F) dikaryon mating in V8 agar. Two independent transformations were performed and indicated by Round #. Shown is one representative experiment.

**Figure 3.**
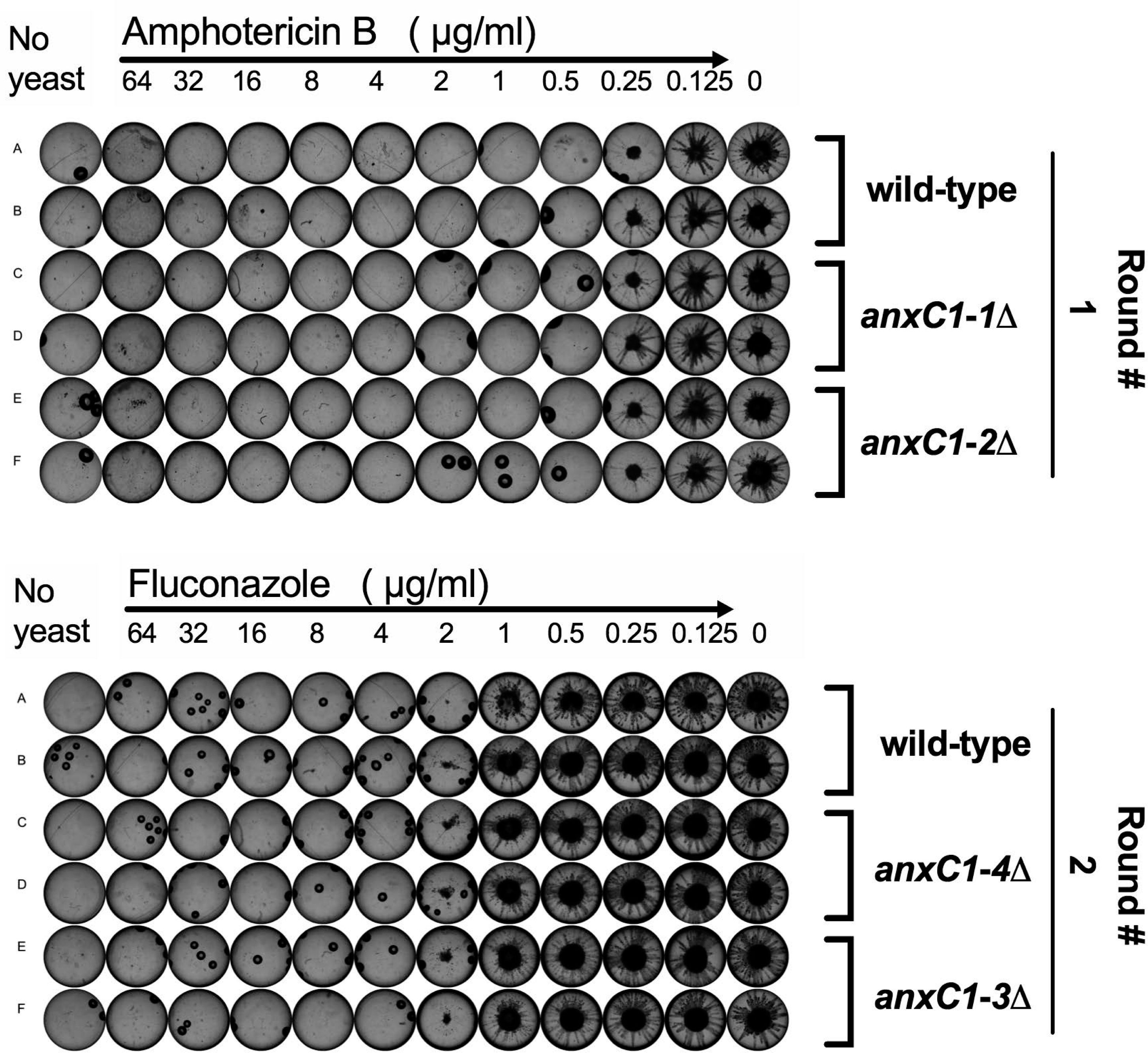
AnnexinC1 deletion does not affect resistance to antifungal drugs. Fungal growth measured by broth microdilution in presence of amphotericin B (top) and fluconazole (bottom). Two independent transformations were performed and indicated by Round #. Experiments were performed three times and shown is one representative experiment.

Fungal annexins are poorly characterized and the literature provides no additional clues to their functions. We performed a broad panel of assays to pinpoint the functions of this protein in fungi (Figure 4). Annexins can be involved in oxidative stress (17). However, we detected no alteration in growth when *anxC1*Δ strains after exposure to oxidative and nitro-oxidative stress. Alternatively, annexins can be involved in membrane trafficking, i.e., exocytosis and plasma membrane dynamics. We reasoned that this could affect secretion of virulence factors and cellular morphology in stress conditions. We found no defect in urease or melanin secretion or alteration in capsule size in our experimental conditions. These results provide evidence against a defect in secretion of virulence factors and therefore a role of cryptococcal annexin in secretion of these virulence factors. However, we found that one of our deletion strains had an increase in cell body size compared to its parental strain. Strikingly, we detected a significant increase in the number of titan cells produced by *anxC1Δ* strains *in vitro* (35) when compared to the parental strain H99, implicating AnxC1 as a possible negative regulator of titan cell formation. We note the magnitude of this effect is small and therefore its biological relevance is difficult to ascertain.

**Figure 4.**
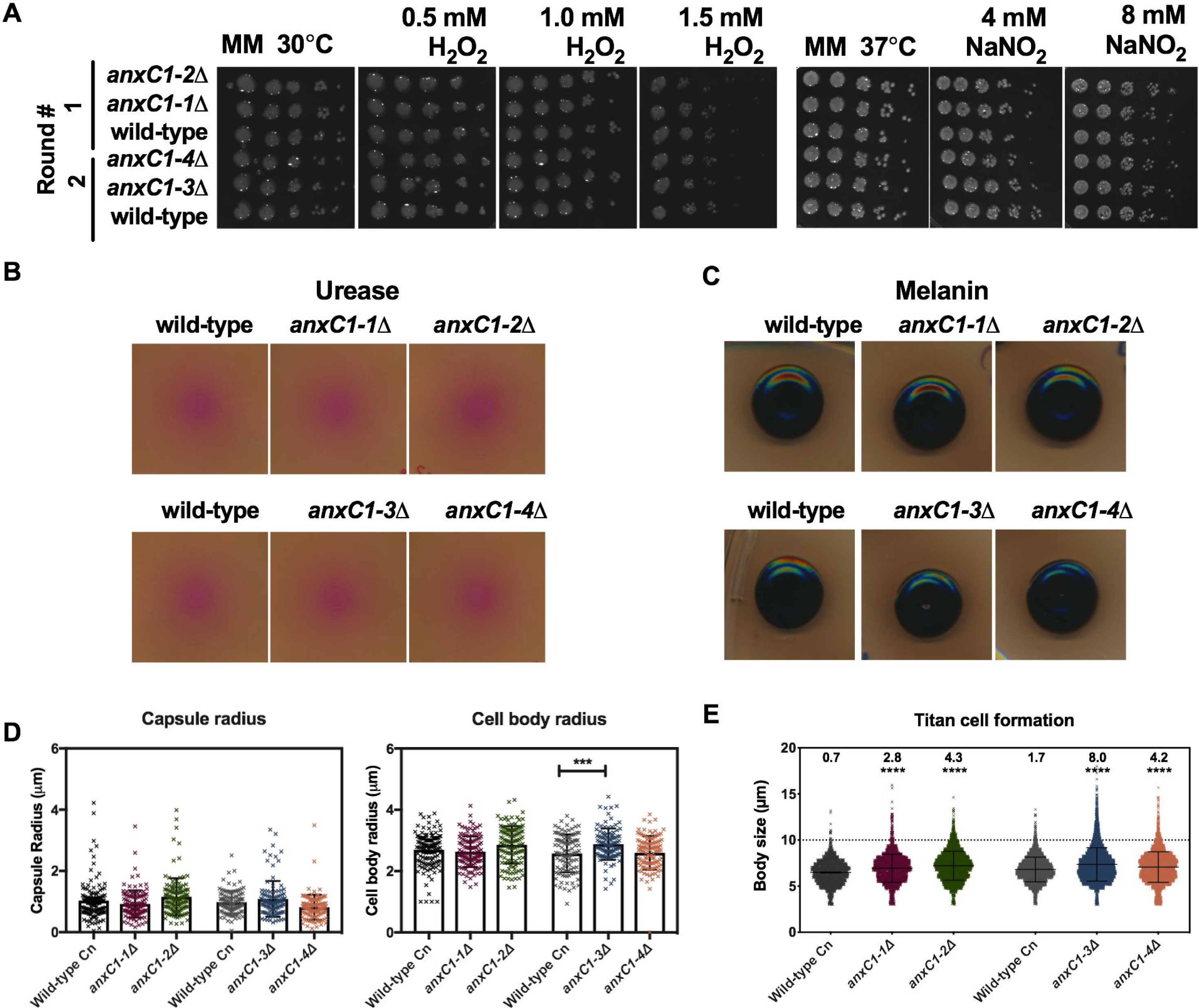
AnnexinC1 deletion does not affect virulence factors but increases production of titan cells. A) resistance to oxidative and nitrosative stress in minimal media agar plates. Two independent transformations were performed and indicated by Round #; B) urease secretion in Christensen agar plates; C) melanin secretion in minimal media with addition of 1 mM L-DOPA; D) capsule size or cell size, after 24h incubation in conditions of mammalian cell culture. ***p<0.001 by one-way ANOVA, with Sidak correction; E) Titan cell production (numbers on top represent % of titan cells). Black bars represent average and SD are shown, **** p<0.0001 for unpaired t-test for all strains compared with parental wild-type strains.

Given the lack of clues to cellular functions of AnxC1 we tested *anxc1Δ* strains for alterations in virulence in several models of virulence that are routinely performed in our laboratory (Figure 5). We found a similar survival rate when we exposed both deletion strains and parental strains to murine macrophages. Because disease is a complex interaction between host and pathogen we decided to measure animal survival when challenged with *anxC1*Δ strains, and we used different routes of infection to cover the broadest range of host-pathogen interactions. Infection of mice via intranasal (IN), intravenous (IV), and intratracheal (IT) routes demonstrated indistinguishable death rate and time to death between *anxC1Δ* strains and their parental wild-type. *C. neoformans* when in environmental niches, soil and trees, is exposed to predation by amoeba. We found that *anxC1*Δ strains had similar survival rates, as compared to wild-type, when ingested by the amoeba *Acanthamoeba castellanii*. When *C. neoformans* is exposed to amoeba in a solid agar matrix, it can filament and form pseudohyphae structures (36)(Fu, manuscript in preparation), but we observed no gross differences between the wild-type and the deletion strains in filamentation morphology (data not shown). Finally, in the *Galleria mellonella* invertebrate infection model, both deletion or wild-type strains showed similar survival kinetics. Based on this data, we conclude that AnxC1 is not required for virulence as measured by in these systems.

**Figure 5.**
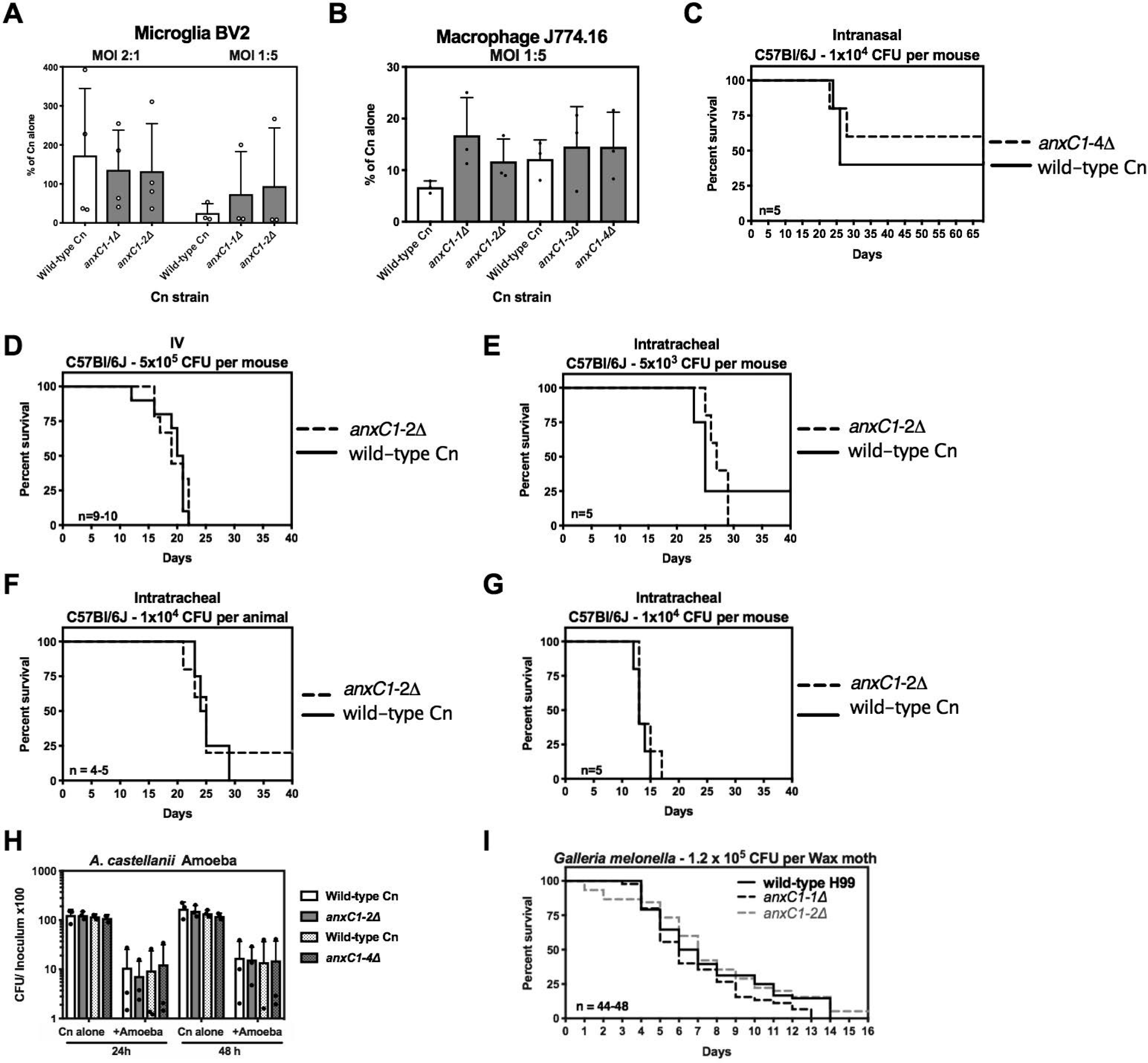
AnnexinC1 deletion does not affect virulence of *C. neoformans*. A) Survival of *C. neoformans* (Cn), as measured by tadpole assay, after 24h infection of BV2, a murine microglia cell line, or B) J774.16, a murine ascites-derived macrophage cell line; C) intranasal infection; D) IV infection; E-G) Intratracheal infections; H) CFU after interactions with *A. castellani*. I) Survival of *Galleria melonella* challenged with Cn strains. Experiments (A-B, H) were repeated three times with triplicates and shown is mean and standard deviation of all experiments. For survival experiments (C-G,I) n represents number of infected individuals per Cn strain and shown is pooled data from all experiments. No differences were found when testing statistical significance by one-way ANOVA (A-B, H), and log-rank or Gehan-Breslow-Wilcoxon test (C-G,I).

## DISCUSSION

The importance of annexins to mammalian and plant biology led us to wonder about the role of fungal annexins, specifically in *C. neoformans.* The scarcity of knowledge regarding fungal annexins and our preliminary searches implicating annexin in responses to two major regulators of fungal virulence, the calcineurin and the Rim101 pathways, encouraged us to pursue a characterization of the cryptococcal annexin. Based on the most recent annexin nomenclature classifications, we propose that the name annexinXIV should be replaced by AnnexinC1.

Our initial search through available datasets suggested that in *C. neoformans*, annexin could be involved in the calcineurin pathway, given that it was previously identified as a target of Crz1 and upregulated in high temperature (22). When testing for phenotypes indicative of deficiencies in calcineurin/Crz1 pathway, AnxC1 is not an effector of the known functions of calcineurin/Crz1. Another master regulator of cryptococcal biology and virulence that was previously shown to affect *ANXC1* expression levels was Rim101, which is critical for cell wall integrity (28), as well as Titan cell induction. Titan cells are enlarged *C. neoformans* cells occurring in mouse lungs during infection and that are generated *in vitro* under multiple conditions (35, 37, 38). The characteristic cell enlargement is associated with virulence and has been shown to be negatively regulated by *Pkr1* (35) and positively regulated by Rim101 (28). The observed increase in titan cell production in *ANXC1*-deletion mutants posits a negative regulation circuit between Rim101 and AnxC1.

We did not observe differences in virulence between *anxC1Δ* and wild type strains in either mammalian or invertebrate infection models. Overall, the lack of phenotypes associated with AnnexinC1 was puzzling, particularly given that its transcription is altered by important virulence regulators such as Crz1 and Rim101. It may be that annexinC1 functions are much subtler than our assays could detect. For example, in the slime mold *D. discoideum*, deletion of annexin delays, but does not prevent, repair of the plasma membrane upon laser wounding (39). An alternative hypothesis is that an additional annexin is present in *C. neoformans* and that both annexins play redundant roles. However, sequence analysis revealed no other annexin in *C. neoformans* and therefore if another molecular player is responsible for redundancy in annexin functions, it could not be readily identified by sequence homology. Our data is similar to what has been previously shown for *Aspergillus* spp, where no obvious phenotypes were observed upon deletion of annexin genes (13, 15, 16). Likewise, in *Neurospora crassa* an annexin deleted strain had no gross growth or filamentation defects (40).

In contrast, in the thermophilic fungi *Thermomyces lanuginosus* a deletion mutant of AnxC7 displayed normal growth in standard media, but had increased resistance to cell-wall stress and oxidative stress. This fungal annexin C7 is predicted to localize to fungal mitochondria (17). These authors also observed that conidia formation was precocious. This observation may be somehow related to the observation that in *C. neoformans* expression of AnxC1 peaked at the time of bud formation (41), approximately 50 min after separation of cells by centrifugal elutriation, insinuating that cryptococcal annexin, and maybe fungal annexins, are involved in membrane dynamics during cellular replication. This framework would fit within our observation that AnxC1 deletion leads to an increase in the formation of titan cells, which display very atypical size and replication patterns, requiring considerable coordination with cellular membrane dynamics. AnxC1 was not differentially regulated during Titan cell formation using Trevijano-Contador et al. dataset (the only RNAseq dataset available to date for Titan cell formation) generated after 7 and 18h of Titan cell induction as compared to normal sized cells (37). A role in membrane dynamics would also fit within the generalized function of annexins as a bridge between phospholipid structures, given the dynamic membrane remodeling needed during cellular replication. Given the scarcity of studies focusing on fungal annexins and shortage of (dramatic) phenotypic insights from deletion mutants, the role of cryptococcal annexins remains cryptic.

In summary, *C. neoformans* possesses one single annexin gene as evident from sequence homology analysis. We were not able to identify a molecular function for AnxC1 because comparisons of annexin-deficient to wild-type strains revealed no discernible phenotype with the notable exception of an increase in titan cell formation. Although it is conceivable that a critical function lurks beneath the absence of vivid phenotypes, we were unable to find a strong effect in major phenotypic features of *C. neoformans* and its virulence in mammalian, invertebrate and protozoal models of infection. In this regard, our experience is similar to that of investigators who have explored the role of annexin in *Neurospora crassa* and *Aspergillus* spp. (12, 13, 15) who have found only weak phenotypes upon deletion of this gene. The explanation for these negative results could lie in the existence of redundant mechanisms to compensate for the absence of annexin or new functions that were not considered in experimental design. In this regard, it is possible that annexins in fungi function very differently than annexins in the other kingdoms of life. Although negative results are disappointing, we take the optimistic view that a gene that is found as a single copy in *C. neoforman*s with homologs throughout the fungal kingdom must be important somewhere in the biology or life cycle of these organisms. Perhaps as new assays are developed to study *C. neoformans* physiology and mechanisms of virulence, a role for its single annexin will become apparent. Accurate three-dimensional structures of fungal annexin proteins, coupled with in-depth cellular localization and transcriptional studies, would allow *in silico* structure homology search, which could help resolve the conundrum of specific biological function of cryptococcal annexins.

## Materials and Methods

### Phylogenetic and protein modelling analysis

Conserved domains were detected with NCBI Conserved Domains (42). Fasta files were downloaded and aligned using MUSCLE (MUSCLE: multiple sequence alignment with high accuracy and high throughput, Nucleic Acids Res., 2004b, vol. 32(pg. 1792-1797) with Mega7 software. This software indicated that phylogenetic analysis should be constructed with Maximum likelihood tree, Neighbor joining analysis, LG+G model with partial deletion. Bootstrap method was performed with 500 replications and phylogeny was found to be reliable between fungi annexins was above 95% (43, 44). The tree is unrooted. A model of AnnexinC1 was built using Swiss-Model workspace (https://swissmodel.expasy.org/interactive#structure) (45, 46), using AnnexinA5 from *Homo sapiens* as a template. FungiDB (http://fungidb.org/fungidb/) was used to look for gene expression patterns of CNAG_02415 (47).

### Cn strains and mutant preparation

The strain H99 wild-type strain was obtained from Jennifer K. Lodge laboratory (available at Fungal Genetic Stock Center) and is also identified as JLCN69, originating from an H99E lineage (Jennifer K. Lodge, personal communication) (48). To confirm some of the phenotypes of mutants, when indicated (in Supplemental Figure 2) we used mutant strains and respective wild-types (one in KN99 and the other in H99 background) from freely available libraries, prepared by the laboratories of Suzanne E Noble and Hiten D Madhani (34). Cells were kept in 10% glycerol frozen stocks. *C. neoformans* mutants were prepared by biolistic transformation (49), using double-joint nourseothricin-split (NAT) markers (50). We joined the NAT markers with sequences flanking 1000-500 bp of CNAG_02415 coding region, which were deposited into gold carrier particles and shot into a lawn of *C. neoformans*. Single colonies were picked from 100 µg/ml NAT plates. We selected several clones and further characterized them via PCR to verify insertion in the correct genomic location, as well as Southern blot using DIG-High Prime DNA labeling and Detection starter kit II (Roche to verify only single insertion of the NAT cassette into each of the strains (577bp). PCR 1 amplifies the entire NAT cassette (Forward primer: 5’-GCCTGATTGATTTGCTGTGG and Reverse primer: GCTTCTCGTTTACAAACAGCGC); PCR 2 is from 994 bp upstream of Annexin gene and 1217 bp internal to *ANXC1* (GCCTGATTGATTTGCTGTGG and GCTTCTCGTTTACAAACAGCGC); PCR 3 is from 1140 upstream of insertion site and 290 bp internal to NAT (forward primer: GTGAGGGAAAGAATTCGTCG and reverse primer: TGTGGATGCTGGCGGAGGATA); PCR 4 is from 847 bp internal to NAT and 65 bp downstream of insertion site (Forward primer: CTCTTGACGACACGGCTTACCGG and Reverse primer: GGTGGACTAAATGGGGTTCAAAGG). Southern blot probe was constructed with primers CTCTTGACGACACGGCTTACCGG and GCCTTCACGAATTCTTAGGGGC recognizing the NAT cassette. Two independent rounds of biolistic transformation, clone screening by PCR and Southern blot were performed and we selected 2 mutants from each round of biolistic transformation, for a total of 4. Mutants are always compared to their parental isolate (from the same round of biolistic transformation).

### Spotting assays

Yeasts were grown overnight at 30°C in YPD broth with shaking. Early stationary phase cultures were diluted to 1×10^6^ cells/ml and then serially diluted 10-fold. The dilutions were spotted (3µl) into each plate and incubated at 30, 37, or 39°C until visible colonies developed. Yeast Peptone Dextrose (YPD) was purchased from DB. Minimal media (MM) consisted of 15 mM glucose, 10 mM MgSO_4_, 29.4 mM KH_2_PO_4_, 13mM glycine, and 3 µM of thiamine-HCl, with final pH=5.5. All chemicals were purchased from Sigma and used in the concentrations indicated in the figure legend. Assays were by incorporating the indicated amount of chemicals in minimal media agar, with the exception of cell wall stress tests which were incorporated into YPD agar plates. Alkaline stress was induced in minimal media supplemented 150mM HEPES and adjusted to indicated pH. Urease secretion was performed in Christensen’s agar and melanin secretion was measured by adding 1 mM L-DOPA to minimal media agar. Dikaryon mating was induced by mixing equal suspensions of alpha and a strains in V8 juice agar (51).

### Antifungal Drug Resistance

The susceptibility of yeast strains to fluconazole and amphotericin B was assessed by broth microdilution according to CLSI M27-A3 (52). Briefly, strains were grown at 35°C for two days on Sabouraud dextrose agar, then used to inoculate drug plates to final concentration of 1.5 x 10^3^ CFU/ml (determined by hemocytometer, rather than spectrophotometer). Plates were incubated for three days at 35°C prior to reading and each well imaged (Cellular Technology). Minimal Inhibitory Concentration (MIC) was determined visually, with fluconazole MIC being the lowest concentration at which growth was at least 50% inhibited, and amphotericin B MIC being the lowest concentration at which no growth was observed. Each strain was tested in duplicate in three independent experiments.

### Macrophage Growth Assays and Capsule Measurements

Two macrophages cell lines were used for most experiments: the macrophage-like murine cell line J774.16 (53) and the microglia-like cell line BV2. The BV2 cell line was a kind gift from Herbert W Virgin. J774.16 were kept in DMEM complete media consisting of DMEM (CellGro), 10% NCTC-109 Gibco medium (LifeTechnologies), 10% heat-inactivated FBS (Atlanta Biologicals), and 1% non-essential amino acids (CellGro). Macrophages were plated at a density of 5 x 10^4^ cells /mL the day before infection. *C. neoformans*, opsonized with 10 µg/ml of mAb 18B7 (54) were added at a Multiplicity of Infection (MOI) indicated in the figure legend in a final volume of 250 µl and infection proceeded for 24h. Macrophages were lysed by resuspending all cells with 10x the volume of distilled water and number of surviving yeasts was measured via a “tadpole” assay (55). Briefly, the aqueous suspension of yeast cells and lysed macrophages is serially 10-fold diluted in YPD broth and allowed to grown overnight at 30°C. Colonies are visible in the wells and the number of surviving yeasts is calculated by multiplying the number of colonies by the dilution of each well. For capsule growth *C. neoformans* cells were incubated in macrophage media in 37°C, 5% CO_2_ for 24h, conditions known to induce capsule growth.

### Amoeba Growth Assays

Amoeba infections were performed as described previously (56). *Acanthamoeba castellanii* strain 30234 was obtained from the American Type Culture Collection (ATCC). Cultures were maintained in PYG broth (ATCC medium 712) at 25°C. *C. neoformans* was grown in Sabouraud dextrose broth with 120 rpm shaking at 30°C overnight (16 h) prior to infection of amoeba. *A. castellanii* cells were washed twice with Dulbecco’s phosphate-buffered saline (DPBS; Corning, Corning, NY), supplemented with Ca^2+^ and Mg^2+^ and 1×10^4^ cells seeded in DPBS into each well of 96-well plates. After 1 h adhesion, 1×10^4^ cells of *C. neoformans* in the same DPBS were added to wells containing amoebae or control wells containing DPBS alone, and the plates were incubated at 25°C. At 0, 24, and 48 h, the amoebae were lysed by repeated shearing through a 27-gauge needles. The lysates were serially diluted, plated on Sabouraud agar, and incubated at room temperature for 48 h for CFU determination. For solid agar matrix infection experiments (pseudohyphae formation), 200 yeast cells were spread on Sabouraud agar, and incubated at 30 °C overnight. *Acanthamoeba castellanii* at a density of 5 x 10^3^ cell per plate were randomly spotted on the agar. The plates were incubated at 25 °C for 2-3 weeks until filamentation was observed macroscopically. Filamention sites were observed for fungal morphology under regular light microscopy.

### Titan Cell Generation

Titan cells were generated according to the protocol generated by Hommel *et al.* (35). Briefly cells were pre-cultured in YPD overnight at 30°C with slow shaking, washed in minimal media and diluted to 1×10^6^ cells/ml in MM. Cell suspension was placed in closed 1.5 ml microcentrifuge tubes and shaken for two days in for two days in Eppendorf Thermomixer^®^ (Hambourg, Germany) at 30°C with 800 rpm shaking. Cells were imaged and cell size (based on cell wall) was determined using ICY software (35). Cells with body size >10 µm were considered Titan cells. Results are expressed as median cell size [interquartile range, IQR] or as median frequency % of titan cells for each strain.

### Galleria Virulence Assays

*Galleria mellonella* larvae were picked based on weight (0.2g ± 0.02g) and appearance (creamy white in color). Larvae were starved overnight at room temperature. Yeast strains were inoculated overnight on Sabouraud broth, 30°C, with shaking. The following day yeasts were washed 3 times with PBS and adjusted to 1.2 x10^7^ cells/ml. Yeasts were injected on the seventh front paw of the larva with 27G tuberculin needles. Infected larvae were incubated at 30°C for 15 days and observed daily for lack of movement (death). Control groups of larvae were inoculated with 10 µL of sterile PBS or left untouched. Experiments were repeated three times with experimental groups of 10-12 larvae at a time (57-60).

### Animal infections

C57BL/6J mice, aged 8–10 weeks, were obtained from Jackson Laboratories and infected intratracheally as previously described with the indicated CFU per animal in a final volume of 50 µl of sterile PBS (61). Intranasal experiments were performed by dropping 40 µl of yeast suspension into the mouse nares while under isoflurane anesthesia (35). Intravenous injections were performed by 40 µl injection in the retroorbital sinus of the animal under isoflurane anesthesia. Mice were monitored daily for signs of stress and deterioration of health throughout the experiment. Animals were euthanized if unable to feed. All animal experiments were approved by Johns Hopkins University IACUC under protocol number MO18H152.

## Acknowledgements

AC was supported by National Institutes of Health (NIH) awards 5R01HL059842, 5R01AI033774, 5R37AI033142, and 5R01AI052733. EF was supported by RISE grant number: 1R25GM113748-01. The fungal deletion libraries were funded by NIH R01AI100272 to Hiten D. Madhani and deposited in the FGSC (http://www.fgsc.net/).

## Author Contributions

CC and MM generated mutant strains and performed the experiments. CC and AC designed the experiments. EC, DSG, MSF, EEF, AA performed the phenotypic assays. CC wrote the manuscript. All authors edited, read and approved the final version of the manuscript.

---

**Supplemental Figure 1.**

Generation of deletion strains (*anxC1Δ*) by double-joint PCR. A) Schematic of deletion cassette; B) Schematic of screening PCR performed; C) PCR and D) Southern blot after digestion with BamHI enzymes for monitoring single insertion of NAT casette. WT represents wild-type parental H99, B represents water blank.

**Supplemental Figure 2.**

AnnexinC1 deletion mutants susceptibility to calcineurin inhibitors, high temperature or CFW-cell wall stress. Strain KN99 and deletion mutant were obtained from the 2015-2016 *Cryptococus neoformans* mutant library (Fungal Genetic Stock Center, 2016 FGSC) and strain H99 and respective deletion mutant were obtained from the 2008 library generated in an H99 background (2008 FGSC) (34). Shown is one representative experiment.

